# Enhancing Recognition and Interpretation of Functional Phenotypic Sequences through Fine-Tuning Pre-Trained Genomic Models

**DOI:** 10.1101/2023.12.05.570173

**Authors:** Duo Du, Fan Zhong, Lei Liu

## Abstract

Decoding high-quality human genomic sequences requires comprehensive analysis of DNA sequence functionality. Through computational and experimental approaches, researchers study the genotype-phenotype relationship and generate important datasets that help unravel complicated genetic blueprints. This study explores the use of deep learning, particularly pre-trained models like DNA_bert_6 and human_gpt2-v1, in interpreting and representing human genome sequences. We meticulously construct multiple datasets linking genotypes and phenotypes to fine-tune pre-trained models for precise DNA sequence classification. Furthermore, we specifically focused on the human endogenous retrovirus (HERV) dataset with commendable classification performance (both binary and multi-classification accuracy and F1 values above 0.935 and 0.888, respectively). We evaluate the influence of sequence length on classification results and analyze the impact of feature extraction in the model’s hidden layers using the HERV dataset. To further understand the phenotype-specific patterns learned by the model, we perform enrichment, pathogenicity and conservation analyzes of specific motifs in the HERV sequence with high average local representation weight (LRAW) scores. Overall, the generated datasets further provide numerous additional genotype-phenotype datasets for evaluating the performance of genomic models. The findings highlight the potential of large models in learning DNA sequence representations, particularly when utilizing the HERV dataset, and provide valuable insights for future research. This work represents an innovative strategy that combines pre-trained model representations with classical omics methods for analyzing the functionality of genome sequences, fostering cross-fertilization between genomics and advanced AI. The source code and data are available at https://github.com/GeorgeBGM/Genome_Fine-Tuning.

## 1. Introduction

The Human Genome Project (HGP) marked the beginning of an era characterized by the assembly to resolve high-quality genome sequences, in which the complete genetic code of DNA is gradually being deciphered[1].The improvement of high-quality reference genome sequences and their gene annotation information, enhancing molecular diagnostics capabilities and enabling advances in disease prevention and personalized treatment strategies [2]. However, the current functional studies on genome sequences focus primarily on ∼3% of protein-coding regions, leaving the vast majority of regulatory functions largely unexplored[3]. Therefore, we need to employ more strategies to apply even the unknown functional regulatory elements in the human genome, such as the recently prominent HERV sequences, which are closely linked with gene regulation, immune modulation, carcinogenesis and the pathophysiology of complex diseases such as neurodegenerative disorders [4].

In the field of artificial intelligence, natural language processing (NLP) has made rapid progress, particularly due to advancements in transfer learning and Transformer architectures, which has led to innovative methods for processing and analyzing large-scale complex datasets [5, 6]. These advances have not only revolutionized traditional text processing tasks, but also provided new tools and methods for fields such as bioinformatics and healthcare. In bioinformatics, the adoption of Transformer models has been particularly transformative in sequence analysis, gene expression, proteomics, and drug discovery, with notable advancements made in the latter two areas[7].

Genomic DNA sequences present unique challenges for machine learning due to their length and complexity. However, their structural similarity to human language (long strings consisting of basic units such as bases or words) provides opportunities for modeling and interpreting DNA sequences using NLP methods[8, 9]. Scientists are increasingly leveraging pre-trained genomic models, leading to significant successes with Transformer-based frameworks [10, 11]and other language framework models[9]. For sequence classification evaluation, Researchers have constructed benchmark datasets for DNA classification and used modified CNN models as the baselines. Ultimately, they found that fine-tuning the pre-trained model DNABERT achieved better performance in classification tasks compared to the DNABERT model with randomly initialized weights and the CNN model[12, 13]. Nevertheless, there is a notable shortage of comprehensive datasets for evaluating the performance of large genomic models, especially for analyses involving complex genotypes[12] [14].

This research has constructed several medically significant genotype-phenotype datasets by utilizing human reference and pan-genome data[15], as well as information from the 1000 Genomes Project. We performed DNA sequence balanced binary classification and imbalanced multi-classification performance evaluation by optimizing pre-trained models such as DNA_bert_6 and human_gpt2-v1. Focusing on the HERV dataset, extensively investigated the representability of the models on this data and identified phenotype-specific motifs within the HERV sequences. Further analysis revealed that genes associated with these motif sequences play critical roles in various biological processes, including neural development and synaptic functions, oncogenesis, cellular adhesion, and spatial localization. The genotype-phenotype dataset we constructed can be used for DNA sequence modeling and performance evaluation of pre-trained large genomic models, providing new insights into the functionality of genomic sequences when exploring the HERV dataset based on fine-tuned models.

## 2. Materials and Methods

### 2.1 Pan-genomic phenotype classification dataset construction

#### 2.1.1 Screening of potential phenotype-related regions

##### 2.1.1.1 HERV Regions

When delineating HERV regions, the initial phase involves acquiring multi-tissue HERV expression annotations as identified by She J et al. [16]. These are merged for adjacent intervals within 10 bp using Bedtools Merge [17]and annotated transcripts are preserved using Bedtools Intersect, ensuring the final interval exceeds 150 bp, which represents the HERV_Coding region. Subsequently, Non-coding HERV regions overlapping with these coding regions are subsequently removed using Bedtools Subtract, yielding HERV_Non-Coding candidate regions.

During the Non-HERV interval selection, genomic intervals from gene annotation files are processed to exclude possible HERV_Coding regions, and contiguous intervals within 10 bp are merged. Non-HERV random sequences with a length distribution similar to HERV_Coding regions are then generated using Bedtools Shuffle, termed Non-HERV_Coding regions. The potential HERV Non-Coding regions are excluded from the non-gene regions of the genome, and using Bedtools Shuffle to generate non-coding region intervals with a length distribution similar to the HERV_Non-Coding region, which is the Non_HERV-Non_Coding region. An assessment phase is then conducted to ensure that there are no overlaps among these four distinct intervals.

##### 2.1.1.2 Immunogenetic Regions

For Immunogenetic regions, immune genes from the ImmPort database[18] are converted to hg38 coordinates and paired with gene annotation insights. Key immune segments identified from the literature are then mapped to their respective genome coordinates. The LRC/KIR region (chr19:54015634-55094318) is divided into ∼10 kb intervals, creating the LRC/KIR immune regions, while the remaining immune zones are also divided into Non-LRC/KIR immune regions.

The selection of Non-Immunogenetic intervals expands all immune regions by ∼1Mb, and complementary genomic regions on hg38 are identified using Bedtools Subtract. Random segments with numerical and length equivalence to all immune region sequences (LRC/KIR and Non-LRC/KIR) are generated within these expanded regions using Bedtools Shuffle, forming Non-Immunogenetic regions. A verification process ensures that these three derived intervals are distinct and do not overlap.

##### 2.1.1.3 Regulatory Regions

The screening of Regulatory Regions involves downloading regulatory features annotation files from Ensembl, from which key regulatory region information (CTCF_binding_sites, Open_chromatin_regions, Enhancers, TF_binding_sites, Promoters) is extracted. To better distinguish the differentiation of specific sequences among various regulatory elements, ensuring the preservation of segments that are at least 150 base pairs long as distinct regions within these elements.

For Non-Regulatory intervals, regulatory elements from the hg38 are excluded. To maintain the same number of regions as in the regulatory elements, we randomly generate equivalent regions within the remaining regions using Bedtools Shuffle. The final check phase confirms that there are no intersections among the six potential intervals.

##### 2.1.1.4 Diseases_GWAS Regions

The selection phase for Diseases_GWAS intervals involved integrating human GWAS signal loci, obtained from UCSC within a genomic coordinate range of 10 kb, with segments predicted by PrimateAI [19]to have a disease score above 0.75. These integrated segments were further classified using the genome annotation file (Gencode.v43) into gene coding and non-gene segments. The segments were further subdivided into 10 kb regions using Bedtools Makewindows, preserving intervals greater than 150 bp for ensuing analyses, designated as Diseases_GWAS_Coding and Diseases_GWAS_Non-Coding regions.

During the selection phase for Non-Diseases_GWAS intervals, Bedtools Shuffle was applied to randomly select Non-Diseases_GWAS_Coding regions with similar number and length distribution as the Diseases_GWAS_Coding regions. A corresponding method was employed for genomic non-gene regions to identify Non-Diseases_GWAS_Non-Coding regions similar to the Diseases_GWAS_Non-Coding regions. Finally, in the interval checking phase, we confirmed that there was no overlap among the four potential regions.

##### 2.1.1.5 Highly_Specifically_Gene Regions

During the selection phase for Highly_Specific_Gene intervals, we noted that olfactory receptor and defensin family genes exhibit distinct tissue-specific protein expression patterns [20, 21]. Consequently, we extracted defensin family genes from the genome annotation file (Gencode.v43), merged intervals within 10 kb using Bedtools Merge, and then divided them into 1 kb sections as Defensins regions with Bedtools Makewindows. Similarly, we followed the same procedure for human olfactory receptor family genes and generated Olfactory_Receptor regions.

For the selection of Non-Highly_Specifically_Gene intervals, we expanded the Highly_ Specifically_Gene intervals by about 1Mb, then extracted the corresponding complementary hg38 genomic regions. By utilizing Bedtools Shuffle, we generated regions with distributions reflecting the Highly_Specific_Gene intervals for both Defensins and Olfactory_Receptor regions. These regions were designated as Other regions. During the checking phase, we verified that the three identified regions did not overlap.

##### 2.1.1.6 Random Regions

In the Random intervals selection phase (selection of 200 million sequences; 10 kb), we distributed the required intervals among the chromosomes according to the hg38 sequence length. Using Bedtools Random, we generated random genomic regions for each chromosome and then merged any overlapping regions using Bedtools Merge. We then calculated the number of intervals on each chromosome, divided them into four files, and randomized the labels to enhance their randomness.

#### 2.1.2 Phenotype Dataset Construction

##### 2.1.2.1 Functional Phenotypic Sequence Extraction within Dataset-Specific Regions

We started using Bedtools Getfasta to extract the regional sequences from the hg38 for potential regions in each dataset. Using Bedtools Intersect, we isolated human pangenomic and 1000G structural variation data [15, 22]. After splitting the continuous sequence with two or more consecutive Ns, retain the part longer than 150 bp and further remove sequences with a similarity above 95% using Mmseqs Easy-linclust [23].

##### 2.1.2.2 Data partitioning for Model Training

For the sequences extracted from each specific region of the dataset, we conducted sequence feature statistics using Seqkit Stat[24]. Then, we employed Seqkit Sample (-s100) to extract approximately 20% of the dataset as an independent test set, while using the remaining approximately 80% as the training and validation set (further divide ∼ 20% as the validation set). For efficient management, we save all data into a single file and provide each dataset with a numerical label (**Supplementary Table 2**).

#### 2.1.3 Display of Phenotype-Related Regions and Datasets

We calculated and plotted the cumulative autosomal sequence length within all phenotype-related areas. Furthermore, we performed statistical analysis to compare the overall distribution and chromosome distribution between sequences in phenotypic and non-phenotypic regions. The gene enrichment rate in specific phenotypic regions was calculated using the following method:

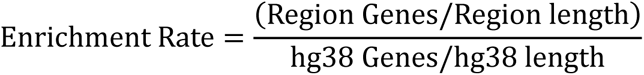

### 2.2 Fine-Tuning of Pre-Trained Models for Phenotypic Datasets

#### 2.2.1 Model Selection and Hyperparameters

##### 2.2.1.1 Overview of core models

The main models, DNA_bert_6 and human_gpt2-v1, are utilized to fine-tune diverse genotype-phenotype datasets, evaluating their ability to classify and recognize functional phenotypic data and assign records precisely to the corresponding labels. DNA_bert_6 processes DNA sequences using 6-mers (stride 1), with pre-training on the BERT architecture, which consists of 12 hidden layers, 12 attention heads, a 768-dimensional hidden layer, and a maximum token limit of 512 for input. The outputs of DNA_bert_6 include attention weights, pooling layers, the final hidden layer, and all hidden layers. On the other hand, Human_gpt2-v1 handles DNA sequences via Byte Pair Encoding (BPE) tokens and is pre-trained with the gpt2 framework on the human telomere-to-telomere genome (T2T-CHM13), which consists of 12 hidden layers, 12 attention heads, a 768-dimensional hidden layer, and a maximum token limit of 1024 for input. The outputs of the model, similar to DNA_bert_6, also include attention weights, the final hidden layer, and all hidden layers.

##### 2.2.1.2 Sequence Classification Model Structure

The pre-trained model can be regarded as data compressor that identifies patterns and latent knowledge within datasets. Therefore, it is possible to incorporate additional network architectures after their hidden layers to achieve sequence classification tasks. In this study, the main approach involved using the AutoModelForSequenceClassification class provided by Huggingface to load pre-trained models (DNA_bert_6 and human_gpt2-v1) for fine-tuning of multiple phenotype datasets. The AutoModelForSequenceClassification class can utilize BertForSequenceClassification and GPT2ForSequenceClassification to apply the pre-trained large BERT and GPT2 models to text classification tasks. The BertForSequenceClassification class adds a dropout layer after the output pooling layer of the pre-trained BERT model and then passes it through a linear classification layer. On the other hand, the GPT2ForSequenceClassification class directly uses a linear classification layer on top of the last hidden layer.

##### 2.2.1.3 Hyperparameters for Training

The fine-tuning hyperparameters for DNA_bert_6 on multiple phenotype datasets are specified as follows: 6-mers (stride 1), batch size of 10, accumulation steps of 4, a learning rate of 2e-5, 50 epochs, and a warmup ratio of 0.1. The human_gpt2-v1 model employs identical hyperparameters for phenotype dataset fine-tuning.

#### 2.2.2 Binary Classification of Functional Phenotypic Sequences

The Fine-tuning for binary classification is balanced across multifunctional phenotype datasets. After processing the sequence data related to phenotype into 6-mers, the first 300 and last 212 6-mer strings in the sequence were extracted. Phenotypic sequences consisting of 512 tokens were then used as the input for the DNA_bert_6 model using padding and truncation strategies. Meanwhile, the phenotype-related sequence data was processed using BPE, and the first 1024 tokens of the sequence were selected using Padding and Truncation strategies as the input for the human_gpt2-v1 model. Subsequently, Independent fine-tuning trials (RUN0, RUN1, RUN2) process this data under specified models and parameters, with evaluations across training, validation, and test sets following a ∼6:2:2 ratio. Various evaluate metrics were calculated after the three rounds of model fine-tuning. Additionally, a voting mechanism was employed, with two identical predicted labels being retained; otherwise, the predicted label from RUN0 (RUN_Vote) was used. The entire training process, facilitated by Huggingface and PyTorch, is streamlined by the Trainer function’s embedded optimization strategies for efficient fine-tuning of large models.

#### 2.2.3 Multiclass Classification of Functional Phenotypic Sequences

The multi-classification of phenotypes was carried out using a strategy similar to binary classification. To attenuate the effects of class imbalance on classification efficiency, the ratio of each data category within the training set labels was calculated and integrated into the CrossEntropyLoss function as a customized Trainer function. Subsequently, the fine-tuning of large models was achieved using Huggingface and PyTorch.

#### 2.2.4 Model Evaluation Metrics

All multiple phenotypic datasets are evaluated using the following metrics while recording the loss and runtime of the model on the test set:

Accuracy is the quotient of the number of correctly predicted samples by the total sample count.

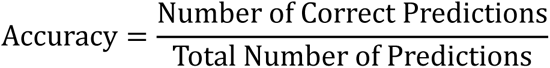

The F1 Score, the harmonic mean of precision and recall, usually used to measure the accuracy of classification models, especially in cases of imbalanced class distribution.

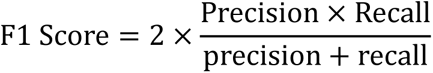

Recall is the ratio of correctly predicted true positives to all actual positives.

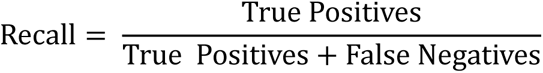

Precision measures the ratio of true positives in all positive predictions.

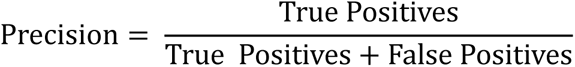

Roc_auc_macro is the area under the ROC curve, used to measure the performance of the model under various classification thresholds and may not accurately reflect the predictive performance of a few classes.

Pr_auc represents the area under the Precision-Recall Curve. Focuses on the predictive power of a small number of classes (positive classes) without being affected by a large number of negative samples, and is suitable for the case of class imbalance.

The MCC (Matthews correlation coefficient) quantifies the performance of binary classification models, considering all varieties of positives and negatives.

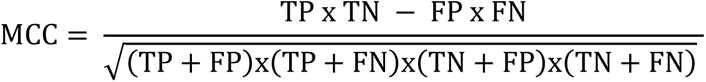

#### 2.2.5 The Impact of Different Lengths of HERV datasets on Phenotypic Classification

To evaluate the classification performance of the models at different sequence lengths, we further evaluate the changes in the prediction results of the DNA_bert_6 and human_gpt2-v1 models within the HERV dataset for three independent fine-tunings (RUN0, RUN1, RUN2 and RUN_Vote). To mitigate the significant differences in sequence length distribution, a logarithmic transformation of the sequence lengths was applied, supplemented by uniformly distributed random noise from -0.1 to 0.1 for length correction. Then, we sort the corrected sequence lengths and select 20 equally spaced values as the interval divisions. For last intervals with less than 10 data points, we merge them into a single interval to maintain reliability and visualization. Next, we present the results of fine-tuning the DNA_bert_6 and human_gpt2-v1 models on the HERV dataset, highlighting the models’ maximum token input threshold and the proportional distribution of data points across intervals.

### 2.3 Model Feature Learning Effect Evaluation in the HERV Dataset

#### 2.3.1 Representativeness of classification labels

DNA sequences in the test set were processed into 6-mer nucleotide fragments. Their frequencies were calculated using CountVectorizer to convert the sequences into numerical feature vectors. The sparse matrices of these feature vectors in a high-dimensional feature space were subjected to dimensionality reduction via TruncatedSVD (n_components=10) and SparsePCA (n_components=10) for further data visualization.

#### 2.3.2 Feature Extraction by the Fine-Tuned Model

The DNA_bert_6 and human_gpt2-v1 models fine-tuned with the HERV dataset were used to extract features in batches from sequences in their test sets. We utilized the final layer of the hidden layer features (batch_size, sequence_length, hidden_size) to calculate the mean feature values across all positions within a sequence, obtaining the representation vector (hidden_size=768) of the entire sequence. A test set feature matrix (sequence_number, 768) was compiled by combining the extracted features from all test set sequences. Subsequently, PCA and t-SNE were applied to reduce dimensions and visualize this feature matrix.

#### 2.3.3 Visualization of HERV Subtype Information

The complete HERV dataset was retrieved from the database, and data from HervD_Atlas[25]were matched with the HERV test set based on hg38 coordinates. The HervD_Atlas fragment with the longest overlap with the test set sequence is selected as the subtype for that sequence. The largest overlapping segment of HervD_Atlas was designated as the subtype for the corresponding sequence. These overlapping sequences were used as reference points in the HERV test set, along with the associated subtype information to interpret t-SNE dimensionality reduction results.

### 2.4 Analysis of phenotype-specific high ALRW in the HERV dataset

#### 2.4.1 Extraction of Attention Matrices from the Fine-Tuned Model using Test Set

Using the DNA_bert_6 model fine-tuned on the HERV dataset, the final layer attention matrices scores (batch_size, num_heads, seq_len, seq_len) for the test set data were extracted. To enhance the representation of the entire 6-mers tokens scores, the attention scores at each position were aggregated with those of the subsequent five positions, resulting in the cumulative score for the entire 6-mers tokens. Given that each position could be included in multiple different 6-mers combinations during cumulative score calculation, it was necessary to record the count frequency at each position to calculate the mean score, which was then normalized across all 6-mers tokens using L2 normalization. Following this approach, we calculated the overall average attention score (Single attentions) for the 12 attention heads, as well as the individual attention scores (Multiple attentions) for each of the 12 attention heads, more accurately capturing the relative significance of each 6-mer Token in the DNA sequence. During the calculation of Single attentions and Multiple attentions for test set data in batches, if the computed length is shorter than 512 tokens, a padded array filled with zeros can be used to avoid errors.

#### 2.4.2 Visualization Attention Scores and Sequence Analysis by Phenotype

Single attentions and Multiple attentions score matrices, derived from the above methods, were combined with the test set labels to examine the distribution patterns of Single and Multi-head attentions across various Token lengths for different phenotypes. To further characterize the sequence feature variations across different Token intervals and explain the DNA sequence reasons for phenotypic Attentions distribution differences, specific Token regions [(0, 50), (50, 150), (150, 250), (250, 350), (350, 450), (450, 512)] were analyzed within the BERT model. The DNA features status in the corresponding token regions for different phenotype sequences ["Non-HERV_Coding", "HERV_Coding", "Non-HERV_Non-Coding", "HERV_Non-Coding"] in the test set were calculated as follows:

GC Content: The proportion of Guanine (G) and Cytosine (C) in the DNA sequence, typically denoted by:

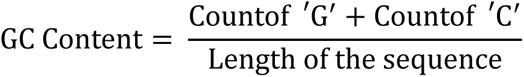

6-mer Frequency: Refers to the number of unique 6-mer sequences in DNA, calculated as:

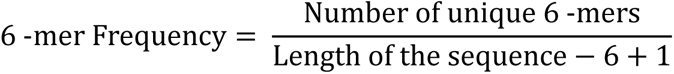

Shannon Entropy: A measure of DNA sequence complexity, computed using the probabilities of each base’s occurrence and applying the concept of information entropy: *H*(*X*) = − ∑ *p*(*x*) ⋅ log2*p*(*x*), where *p(x)* is the probability of the occurrence of base *x*.

CpG Island Score: The observed-to-expected frequency ratio of CpG dinucleotides, calculated based on the occurrence probabilities of C and G within the DNA sequence:

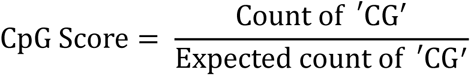

where the expected count of ’CG’ is the product of the frequency of ’C’, the frequency of ’G’, and the sequence length.

Line graphs depicting the variation of DNA sequence features in the different Token regions [(0, 50), (50, 150), (150, 250), (250, 350), (350, 450), (450, 512)] were plotted.

#### 2.4.3 Analysis of Phenotype-Specific HERV Subtypes in the Dataset

In the HERV test set, data that overlapped with the HervD_Atlas database were selected to calculate the percentage of various HERV classes (ERV1; ERV2; ERV3; ERVL-MaLR; Gypsy; LTR) within coding and non-coding regions.

The data were further analyzed to determine the representation of different HERV functional groups. The relative enrichment rates were calculated as the ratio of each group’s percentage in non-coding regions to its percentage in coding regions. This ratio allows us to understand the tendency of each group for being in non-coding versus coding regions, thereby providing insight into the functional dynamics of HERV elements within the genome.

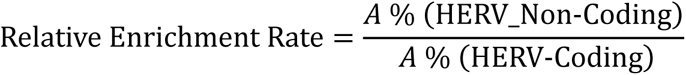

Here, A denotes an individual element within the HERV functional group.

### 2.5 Motifs Analysis using high ALRW Scores in the HERV Dataset

#### 2.5.1 Enrichment of Motifs in continuously high ALRW score sequences

##### 2.5.1.1 Extraction of Continuously high ALRW Score Sequences

In order to extract the continuous high attention score regions in specific phenotypic DNA sequences, the regions with scores exceeding the average value and exceeding the minimum value by 10 times are initially considered as potential areas of interest. These regions are represented as Boolean arrays, with the high attention parts set to true. Then, consecutive true regions with a length greater than 5 bp are identified from the Boolean array as continuous high attention score regions. We then aligned the token coordinates of these segments to the corresponding DNA sequences, extending upstream and downstream by 4 bp each to capture maximum potential information.

##### 2.5.1.2 Motif Analysis of high ALRW Scoring Sequence

DNA sequences from the HERV dataset associated with the identified high attention score regions across different phenotypes ["Non-HERV_Coding", "HERV_Coding", "Non-HERV_Non-Coding", "HERV_Non-Coding"] underwent motifs analysis using Fimo software in conjunction with non-redundant transcription factor binding sites sourced from the JASPAR database in both MEME and TRANSFAC formats. Motifs sequences identified in the above four phenotypes were combined and filtered to retain those with q.value <= 0.05 for further analysis. Firstly, Venn diagrams are used to represent the number intersections of identified motifs among different phenotypes. Secondly, the specificity enrichment of motifs sequences in the HERV group ["HERV_Coding", "HERV_Non-Coding"] relative to the Non-HERV group ["Non-HERV_Coding", "Non-HERV_Non-Coding"] is statistically analyzed. The frequency of motifs sequences occurring in the HERV and Non-HERV groups is normalized by dividing by the cumulative number of motifs sequences in each group. Then, the hypergeometric distribution test is used to perform enrichment analysis on specific motifs, with the calculation method being phyper(*k*-1, *m*, total_motifs-*m*, total_case, lower.tail = FALSE), where *k* is the normalized frequency of each specific motif in the HERV group, *m* is the normalized frequency of each specific motif in the HERV and Non-HERV groups, total_motifs is the total number of motifs in the HERV and Non-HERV groups, and total_case is the total number of motifs in the HERV group. Next, the Benjamini-Hochberg method is used to correct the *p*-values. Based on the frequency of enriched motifs in HERV_Coding and HERV_Non-Coding facilitated categorization of motifs into three categories ["HERV_NonCoding", "HERV_Coding", "HERV_Both"]. Selected motifs sequences are shown for enrichment results (P_hyper_adjust <= 0.05; HERV_Rate >= 0.5; HERV_Num > 10, where P_hyper_adjust is the adjusted *p*-value, HERV_Rate is the proportion of the specific motif’s normalized frequency in the HERV group, and HERV_Num is the normalized frequency of the specific motif in the HERV group).

##### 2.5.1.3 Motifs Analysis of the full-length HERV Dataset Sequences

To validate the reliability of motifs sequences identified by high ALRW scores, we performed a similar statistical analysis and evaluation on the full-length HERV test set DNA sequences as above. We then calculated the differential enrichment of motifs between the two sets and compared them based on the three categories of enriched motifs ["HERV_NonCoding", "HERV_Coding", "HERV_Both"].

#### 2.5.2 Functional Enrichment and Pathogenicity of Phenotype-Specific Motifs

Motifs were matched to the hg38 reference, combining adjacent intervals within a 10 bp range to define HERV sequence-specific motif regions.Firstly, By integrating hg38 gene and regulatory element annotations, we quantified functional element ratios within these specific motif regions. Subsequently, protein-coding genes that overlap with this specific motifs region were identified for functional enrichment and disease correlation analyses via Metascape. Additionally, we compared regions characterized by continuous high attention score motifs with those identified by full-sequence analysis, pinpointing common motifs and those unique to each method. Pathogenicity scores for variants in these regions were predicted using PrimateAI and AlphaMissense[26], and checked for intersection with HervD_Atlas. Suitable examples were selected for display using the UCSC genome browser.

#### 2.5.3 Species Conservation of HERV Sequence-Specificallhy Enriched Motif Sequences

##### 2.5.3.1 Polymorphism in Specific Motif Regions of the Human Pangenome

The genomic diversity within HERV sequence-specific enriched motif regions of the human pangenome was assessed using Odgi Depth [27].Gene annotations that overlapped with these regions were categorized by chromosome and gene category using Bedtools Intersect. We identified genes that exhibited both high divergence and high conservation in the human population and discussed their conservation in primates.

##### 2.5.3.2 Conservation of Specific Motifs in Primates

Genome-wide alignment data from 27 primate species, obtained from UCSC with human hg38 as the reference, were extracted using MafSpeciesSubset. The primate evolutionary tree was reconstructed with PhyloFit, facilitating the evaluation of primate conservation scores using PhyloP and PhastCons. The conservation scores within motif-enriched regions were determined using BigWigAverageOverBed, complemented by gene annotations from Bedtools Intersect to present a comprehensive conservation landscape and annotate specific genes [28].

## 3. Results

### 3.1 Phenotypic Regions Collection and Dataset Construction

#### 3.1.1 Functional Phenotypic Regions of the Human Genome

We consolidated five biomedically significant phenotypes from existing knowledge datasets: Potential Disease Susceptible Regions (Diseases_GWAS), Regulatory Element Regions (Regulatory), Endogenous Viral Regions (HERV), Immunogenetic Regions (Immunogenetic), and Highly Specifically Expressed Gene Regions (Highly_Specifically_Gene). By organizing the genome coordinates corresponding to these phenotypes, we collected essential information about the specific sequences associated with the each phenotypes (**Supplementary Table 1**). For instance, the intervals within HERV dataset cover nearly 302.9 Mb on the human genome, with the longest interval length exceeding 153 kb. There are a total of 382,317 intervals within the genome, covering approximately 98.47% of the HERV intervals from the HervD_Atlas dataset, and spanning a total of 29.68Mb bases [25]. Using the genomic coordinates of these five phenotypes, we illustrated the cumulative length distribution of the phenotype sequence regions across different chromosomes (Fig.1A). The cumulative distributions of Diseases_GWAS and Regulatory Regions are relatively consistent across different chromosome lengths, whereas HERV, Immunogenetic and Highly_Speci fically_Gene show distinct chromosomal enrichments. Notably, Highly_ Specifically_Gene displays extensive enrichment on chromosome 11, primarily due to the predominant distribution of OR (olfactory receptor) genes on this chromosome.To further emphasize the genes as critical functional units, the study also exhibited gene enrichment ratios within the phenotypic regions (Fig.1B), highlighting an enrichment in the number of genes compared to the genome-wide average, particularly for protein-coding genes.

**Fig. 1.**
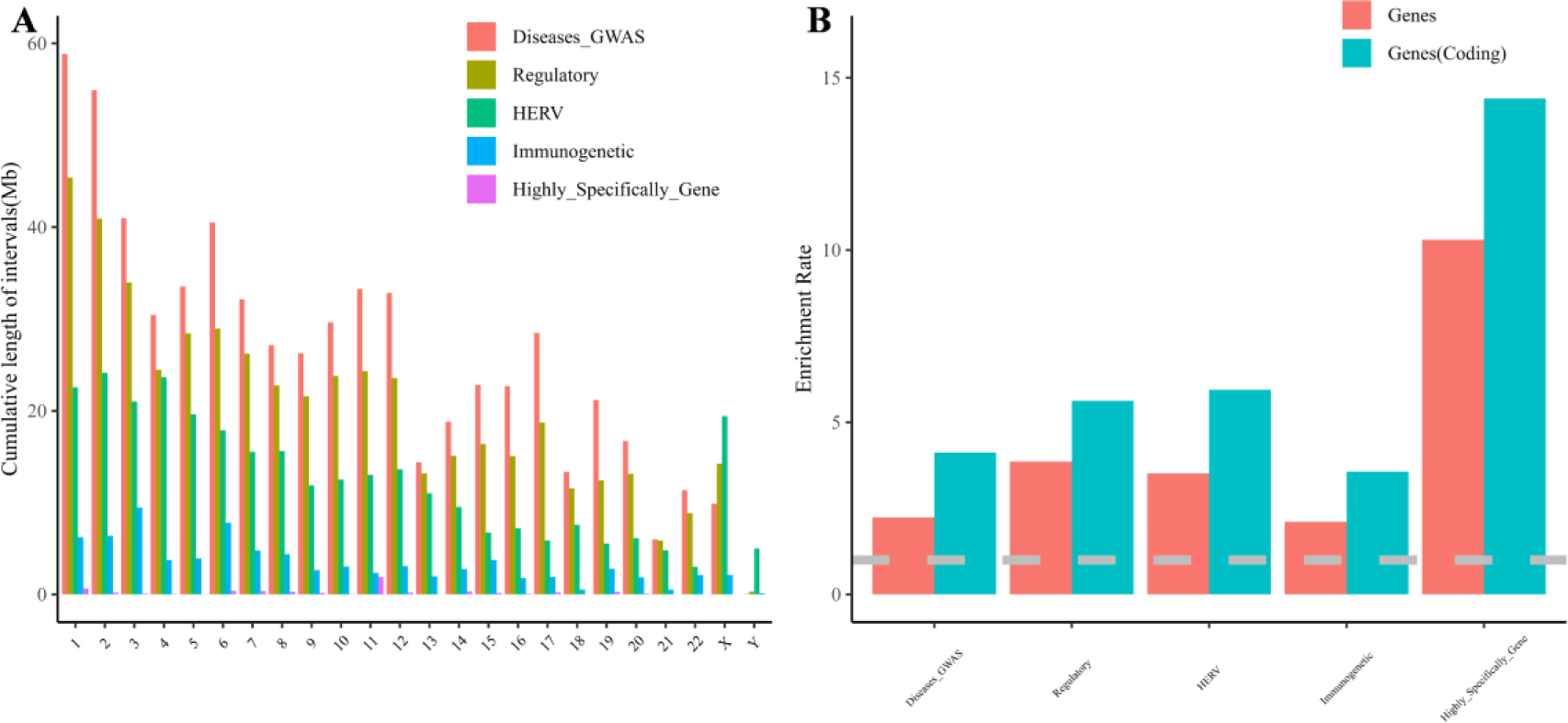
illustrates the distribution and genetic characteristics of the selected phenotypic regions. **(A)** The cumulative chromosomal distribution patterns of five regions with phenotypic data. **(B)** The enrichment rates of all genes (red) and coding genes (blue) within the five phenotypic regions compared to the entire genome. A gray color (y=1) or below indicates the absence of enrichment.

#### 3.1.2 Construction of a Multi-phenotype Classification Dataset for Humans

We constructed multiple functional phenotypic classification datasets by linking specific interval features of genomic phenotypic sequences with randomly selected non-phenotypic genomic regions as controls. We also demonstrated the length distribution of functional and non-functional random regions for different phenotypes (**Supplementary Fig.1A**). The results revealed similar length distributions of these two regions across all datasets. The HERV and Regulatory phenotype datasets, maintaining the original interval lengths, allowed us to analyze the chromosomal distribution of the corresponding functional and non-functional random regions, confirming the uniformity of the constructed datasets across all chromosomes (**Supplementary Fig.1B**).

Subsequently, we extracted corresponding sequences from the selected specific functional and non-functional random regions, using the reference genome hg38, the human pangenome and the 1000 Genomes. After removing redundancy, we performed sequence statistics on the constructed functional phenotype datasets. Furthermore, we classified these datasets into binary and multi-classification categories considering biological factors (Table 1). The HERV dataset, for example, includes both balanced binary classification (HERV: Non-HERV = 417,416: 362,907) and imbalanced multi-classification (HERV_Coding: Non-HERV_Coding: HERV_Non-Coding: Non-HERV_Non-Coding = 16,556: 18,994: 400,860: 343,913). This dataset features approximately 22.64Mb of coding and roughly 295.69Mb of non-coding sequences within the HERV functional regions, allowing for the creation of a multi-classification task using binary classification. Furthermore, the Dataset_Regulatory is the most diverse genotype-phenotype dataset, requiring a six-class classification task to distinguish TF_binding_site, Enhancer, CTCF_binding_site, Promoter, Open_chromatin_ region, and Non-Regulatory element sequences within the genome.

**Table 1.** Display of functional phenotypic dataset sequence.

| Dataset | Classification<br>(Multiclass/Binary) | Numbers | Length<br>(Sum) | Length<br>(Average) | Length<br>(Max) |
| --- | --- | --- | --- | --- | --- |
| Dataset_HERV<br>(HERV) | HERV_Coding/HERV | 16,556 | 22,643,342 | 1,368 | 125,188 |
|  | HERV_Non-Coding/HERV | 400,860 | 295,693,006 | 738 | 112,801 |
|  | Non-HERV_Coding/ Non-HERV | 18,994 | 23,490,411 | 1,237 | 83,426 |
|  | Non-HERV_Non-Coding/ Non-HERV | 343,913 | 251,939,874 | 733 | 93,893 |
| Dataset_Immuno<br>(Immunogenetics) | Immuno_KIR/Immuno | 1,723 | 3,096,500 | 1,797 | 39,269 |
|  | Immuno_Others/Immuno | 16,719 | 83,307,498 | 4,983 | 71,249 |
|  | Non-Immuno | 18,326 | 78,935,043 | 4,307 | 93,519 |
| Dataset_Regulatory<br>(Regulatory) | TF_binding_site/Regulatory | 22,860 | 13,391,832 | 586 | 93,893 |
|  | Enhancer/Regulatory | 320,389 | 335,938,843 | 1,049 | 93,893 |
|  | CTCF_binding_site/Regulatory | 94,738 | 52,273,466 | 552 | 93,893 |
|  | Promoter/Regulatory | 52,413 | 79,591,023 | 1,519 | 99,371 |
|  | Open_chromatin_region/Regulatory | 130,006 | 54,838,694 | 422 | 93,893 |
|  | Non-Regulatory | 494,389 | 435,738,664 | 881 | 96,745 |
| Dataset_Diseases_GWAS<br>(GWAS_loci/PrimateAI-3D scores) | Diseases-GWAS_Coding/Diseases_GWAS | 175,920 | 566,540,577 | 3,220 | 99,371 |
|  | Diseases-GWAS_Non-Coding/Diseases_GWAS | 51,427 | 132,677,910 | 2,580 | 92,731 |
|  | Non_Diseases-GWAS_Coding/Non_Diseases_GWAS | 148,235 | 550,157,564 | 3,711 | 93,893 |
|  | Non_Diseases-GWAS_Non-Coding/Non_Diseases_GWAS | 31,298 | 104,453,319 | 3,337 | 93,893 |
| Dataset_Highly_Specifically_Gene<br>(Defensins/Olfactory Receptor) | Defensins/Highly_Specifically_Gene | 415 | 418,877 | 1,009 | 9,191 |
|  | Olfactory_Receptor/ Highly_Specifically_Gene | 5,400 | 5,305,444 | 983 | 40,645 |
|  | Others | 5,610 | 6,653,032 | 1,186 | 94,155 |

### 3.2 Binary Classification of Functional Phenotype Datasets

#### 3.2.1 Binary Classification Performance of Multiple Phenotype Datasets

To evaluate the performance of fine-tuned pre-training models on multiple phenotype datasets, we conducted three independent fine-tuning iterations (RUN0; RUN1; RUN2) using the DNA_bert_6 and human_gpt2-v1 models for each phenotype dataset, including both training and validation sets. The model performance was then evaluated on the test set using various metrics for all datasets (Table 2), using the random genomic region dataset (Dataset_Random) serving as a foundational baseline. The constructed multiple phenotype datasets demonstrated differential classification performance, with Dataset_HERV, Dataset_ Diseases-GWAS, and Dataset_Immuno models performed well for all metrics, especially Dataset_HERV with accuracy and F1 values above 0.935. In comparison, the DNA_bert_6 and human_gpt2-v1 models showed negligible performance variance, although with a notable increase in fine-tuning time for human_gpt2-v1. In summary, the models’ performance across multiple phenotypic datasets highlights that the phenotypic labels assigned to specific DNA sequences can partly reflect the information contained in the data itself, implying distinctive DNA sequence patterns associated with the HERV, Diseases-GWAS, and Immuno phenotypes. Moreover, the Dataset_Random outcomes imply that current pre-trained models struggle in distinguishing genomic regions in a non-selective manner. This discrepancy likely arises from the incongruence between the information contained in the sequence data and the assigned labels.

**Table 2.** Binary classification performance of functional phenotypic datasets using fine-turned models.

|  | Model | Acc | F1 | Loss | Precision | Recall | Auroc | Pr_auc | MCC | Time |
| --- | --- | --- | --- | --- | --- | --- | --- | --- | --- | --- |
| Dataset_HERV | A | 0.9354±0.0025 | 0.9386±0.0025 | 0.1748±0.0026 | 0.9544±0.0027 | 0.9234±0.0051 | 0.9851±0.0007 | 0.9883±0.0006 | 0.8710±0.0048 | 13.2393±0.3802 |
|  | B | 0.9363±0.0010 | 0.9395±0.0010 | 0.1554±0.0012 | 0.9566±0.0035 | 0.9230±0.0040 | 0.9859±0.0003 | 0.9888±0.0003 | 0.8731±0.0019 | 17.7381±1.2064 |
| Dataset_Immuno | A | 0.8122±0.0002 | 0.8051±0.0037 | 0.3211±0.0013 | 0.8414±0.0132 | 0.7729±0.0182 | 0.9238±0.0007 | 0.9298±0.0007 | 0.6274±0.0017 | 0.7026±0.0165 |
|  | B | 0.8129±0.0008 | 0.8128±0.0039 | 0.3198±0.0021 | 0.8182±0.0167 | 0.8095±0.0244 | 0.9238±0.0013 | 0.9300±0.0011 | 0.6274±0.0018 | 1.9810±0.0044 |
| Dataset_Regulatory | A | 0.7438±0.0003 | 0.7764±0.0028 | 0.5016±0.0013 | 0.7540±0.0060 | 0.8005±0.0127 | 0.8270±0.0005 | 0.8602±0.0004 | 0.4787±0.0009 | 18.4481±0.0137 |
|  | B | 0.7614±0.0012 | 0.7911±0.0020 | 0.4744±0.0015 | 0.7704±0.0061 | 0.8134±0.0104 | 0.8462±0.0014 | 0.8763±0.0012 | 0.5147±0.0027 | 27.2533±1.6940 |
| Dataset_Diseases_GWAS | A | 0.8380±0.0007 | 0.8537±0.0021 | 0.3023±0.0023 | 0.8642±0.0095 | 0.8442±0.0130 | 0.9368±0.0008 | 0.9541±0.0006 | 0.6731±0.0022 | 7.2913±0.0123 |
|  | B | 0.8443±0.0010 | 0.8609±0.0014 | 0.2976±0.0019 | 0.8622±0.0048 | 0.8597±0.0070 | 0.9393±0.0008 | 0.9555±0.0006 | 0.6843±0.0018 | 21.3534±0.0286 |
| Dataset_Highly_Specifically_Gene | A | 0.6346±0.0044 | 0.6081±0.0180 | 0.6075±0.0029 | 0.6688±0.0081 | 0.5604±0.0352 | 0.7076±0.0034 | 0.7372±0.0043 | 0.2758±0.0054 | 0.2051±0.0003 |
|  | B | 0.6237±0.0056 | 0.5861±0.0147 | 0.6349±0.0040 | 0.6668±0.0189 | 0.5264±0.0332 | 0.6894±0.0011 | 0.7261±0.0011 | 0.2572±0.0144 | 0.1858±0.0098 |
| Dataset_Random | A | 0.5205±0.0031 | 0.6503±0.0118 | 0.6953±0.0008 | 0.5299±0.0006 | 0.8436±0.0384 | 0.4983±0.0021 | 0.5296±0.0006 | -0.0003±0.0026 | 0.7690±0.0038 |
|  | B | 0.5134±0.0059 | 0.5895±0.0242 | 0.7010±0.0014 | 0.5327±0.0019 | 0.6656±0.0606 | 0.4997±0.0012 | 0.5290±0.0014 | 0.0085±0.0067 | 2.4468±0.0038 |

*Note:* The **A** represents the DNA\_bert\_6 model; **B** represents the human\_gpt2-v1 model.

#### 3.2.2 Binary Classification Performance Across HERV Sequence Lengths

When fine-tuning the large pre-trained models DNA_bert_6 and human_gpt2-v1 with new datasets, it is necessary to adjust the length of the input DNA sequences to keep them within the maximum allowed tokens for each respective model. Similar strategies have also been applied in the fine-tuning of multiple functional phenotypic datasets, including the HERV dataset. Considering the exceptional performance of the model on the HERV dataset and the preservation of the original interval length of the functional phenotype within this dataset, we investigated the association between the model performance and the sequence length on the HERV dataset. The test set was partitioned into multiple subsets based on sequence length, and the changes in HERV sequence classification scoring metrics within different length ranges were evaluated in three independent fine-tuning experiments (Fig.2). The fine-tuning experiments on the HERV dataset, utilizing the DNA_bert_6 model (512 tokens; Fig.2A) and the human_gpt2-v1 model (1024 tokens; Fig.2B), reveal that as the sequence length surpasses the maximum token limit of the model (red line), the model’s classification metrics gradually decrease. However, owing to the majority of test sequences falling within the token boundaries (under gray line), the overall classification performance remained high. Furthermore, despite the human_gpt2-v1 model having an increased input length compared to the DNA_bert_6 model, the improvement in overall metrics is minimal. Therefore, it is essential to carefully evaluate the data distribution and comprehensively consider model specifications to efficiently optimize task performance.

**Fig. 2.**
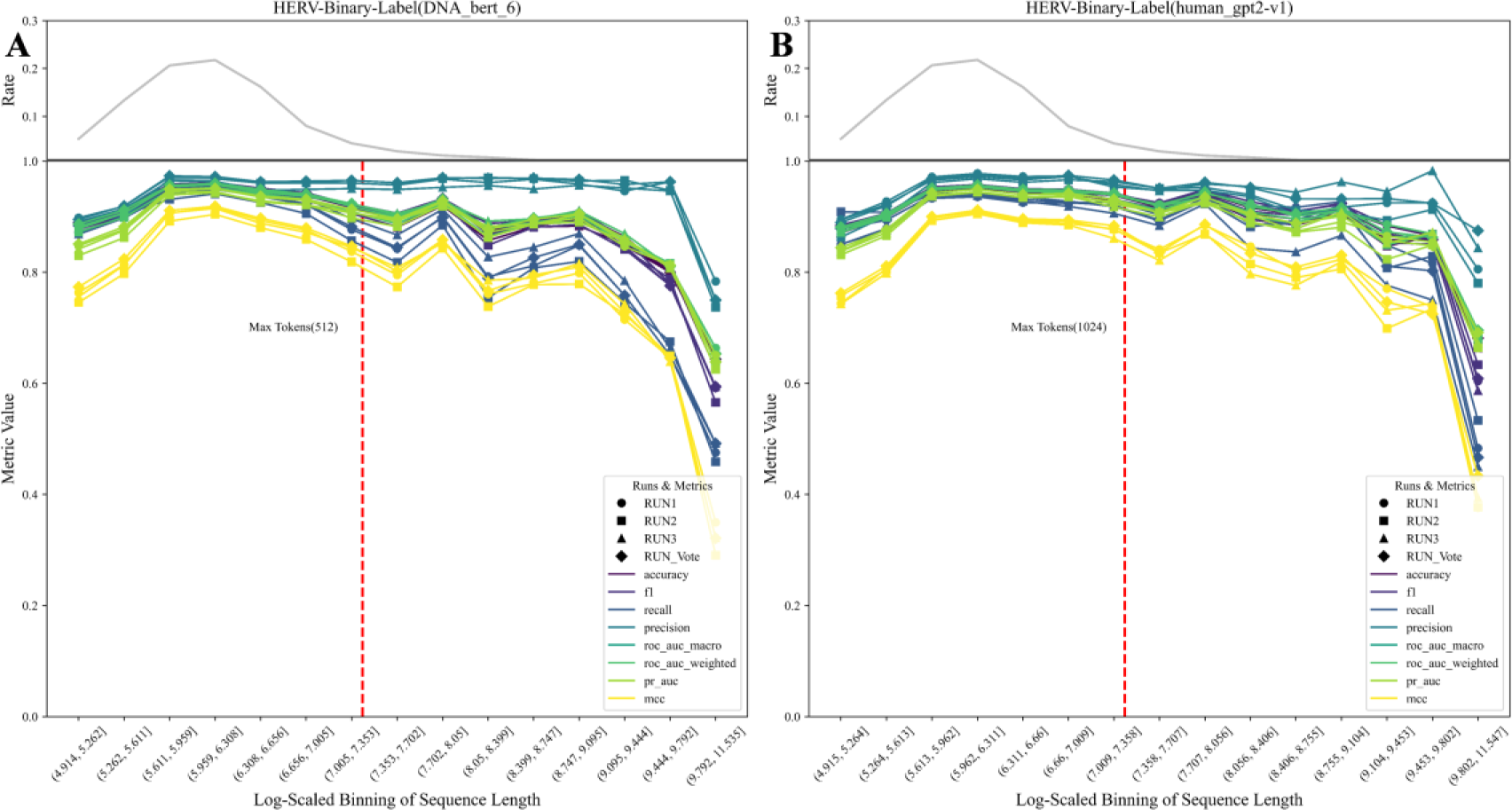
Binary classification effects of sequences across different length ranges in the HERV dataset. **(A)** DNA_bert_6 model fine-tuning results; **(B)** Results of fine-tuning the human_gpt2-v1 model. The upper grey line is the percentage of sequences within different length ranges, the red line represents the maximum tokens allowed by the model, and the RUN_Vote represents the aggregated outcome of three independent voting runs (RUN0; RUN1; RUN2).

### 3.3 Multi-classification of Functional Phenotype Datasets

#### 3.3.1 Multi-classification Performance of Phenotypic Datasets

To further evaluate the multi-classification performance of pre-trained large models on various phenotype datasets, we employed a training and evaluation strategy used in binary phenotypes to evaluate the multi-classification effect. Given the significant class imbalance in certain multiclass phenotype datasets (Table 1), we corrected the cross-entropy loss function of the pre-trained large models DNA_bert_6 and human_gpt2-v1 during the fine-tuning process by adjusting the weights of the labels in the training set. Moreover, we performed three independent fine-tunings rounds (RUN0; RUN1; RUN2) for each dataset, including both training and validation sets, and subsequently evaluated the classification outcomes using the test set (Table 3). Notably, when comparing the multi-classification fine-tuning of the model to binary classification within the same phenotype dataset, we observed a decrease in metrics, which may be attributed to the increased difficulty of the multi-classification task. Consistently, the model’s performance exceeded that of the genomic random region dataset (Dataset_Random) in Dataset_HERV, Dataset_Immuno, and Dataset_Disease-GWAS, with Dataset_HERV achieving accuracy and F1 scores above 0.888. In specific functional phenotypic tasks, the significant decrease in performance between multi-classification and binary classification tasks was observed in Dataset_Regulatory and Dataset_Diseases-GWAS, indicating that too many class labels and inconsistencies between data and label information can adversely affect the final sequence classification performance.

**Table 3.**
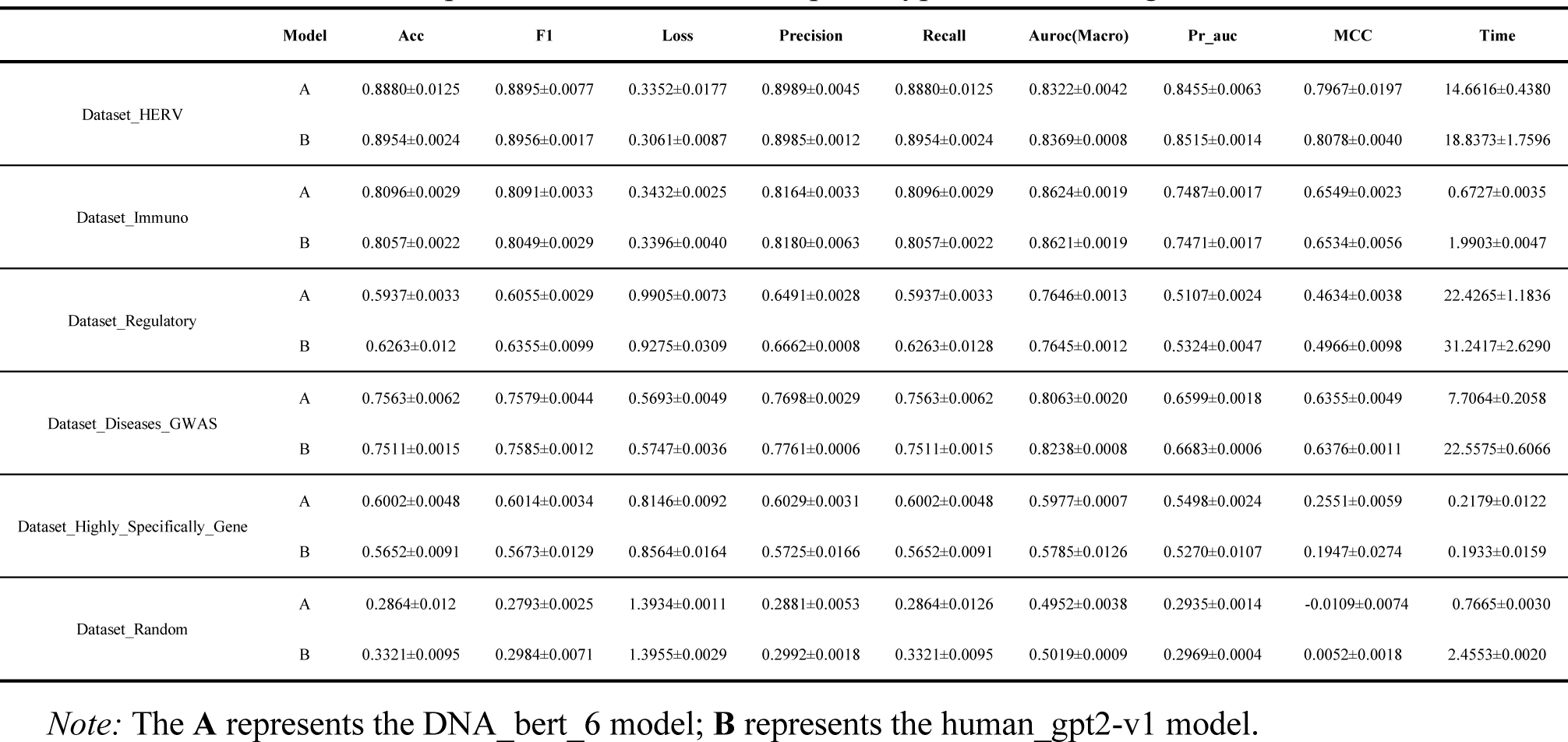
Multi-classification performance of functional phenotypic datasets using fine-turned models.

#### 3.3.2 Multi-classification Performance Across HERV Sequence Lengths

We adopted a similar evaluation approach to that employed in assessing binary classification performance at varying sequence lengths. This study focuses on presenting the multiclass performance of the pre-trained large models, DNA_bert_6 and human_gpt2-v1, on sequences of different lengths within the Dataset_HERV dataset (Fig.3). Similar to its impact on binary classification, increasing the sequence lengths resulted in a decline in multiclass classification metrics, with a notable decrement beyond the model’s maximum Token allowance. The trend observed in multiclass evaluation metrics is consistent with the change in sequence length frequency (gray), this could be attributed to the decrease in the number of test samples for each class in the multiclass task, thereby leading to a significant metrics decrease. The human_gpt2-v1 model (Fig.3B) slightly outperformed DNA_bert_6 (Fig.3A) in handling longer sequences within a permissible range. Nevertheless, considering that the majority of test sequences adhere to the token limit of the DNA_bert_6 model, the differences in the final evaluation metrics are not statistically significant. Moreover, a voting strategy across classification trials may improve model consistency (RUN_Vote), in split of a decrease in MCC metric was observed in RUN1. Therefore, it is necessary to consider the distribution characteristics of the data and classification task complexity comprehensively. Conducting independent multiple experiments to evaluate the overall classification performance of the model is crucial in strengthening its robustness.

**Fig. 3.**
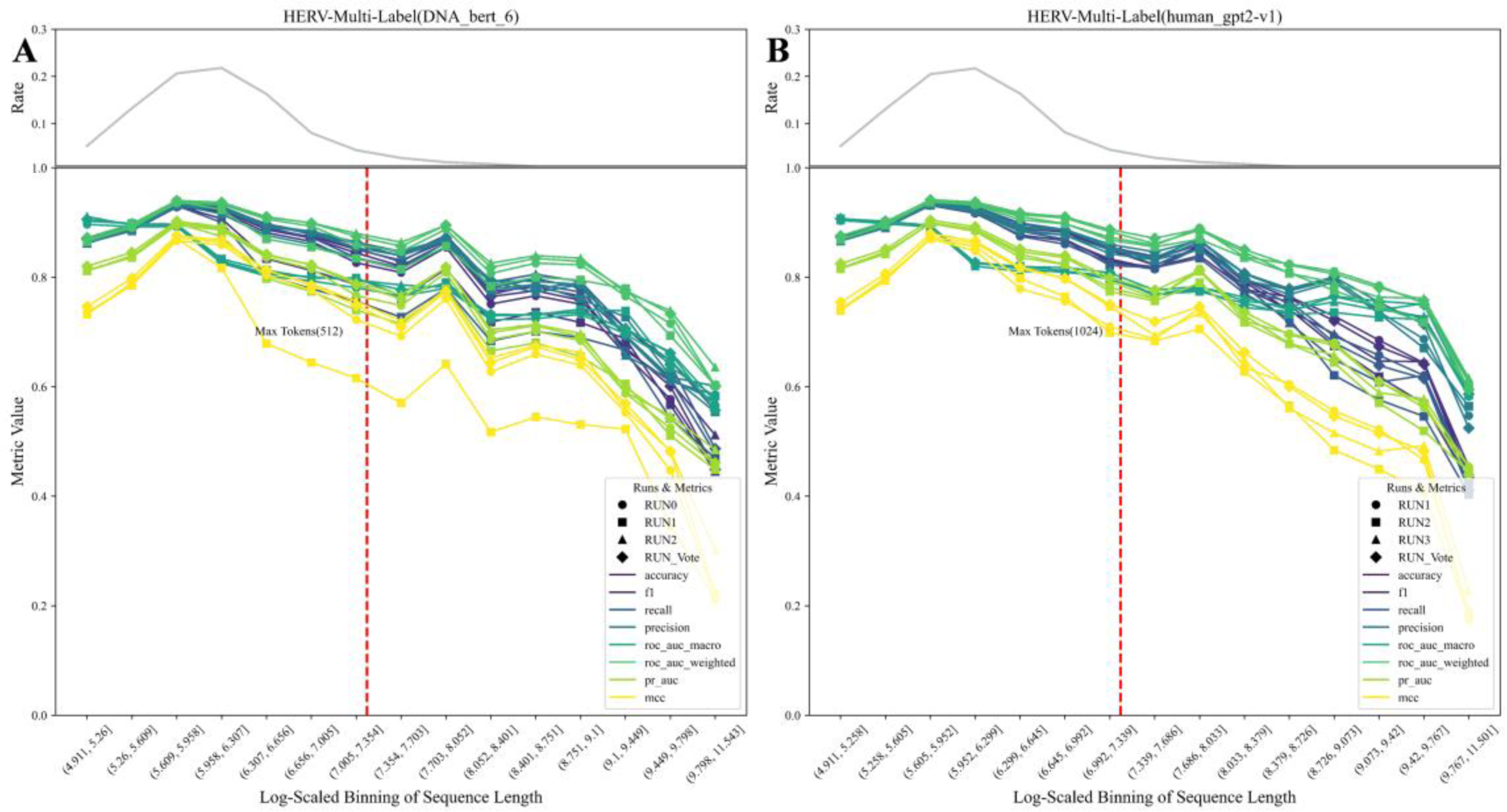
Multi-classification effects of sequences across different length ranges in the HERV dataset. **(A)** DNA_bert_6 model fine-tuning results; **(B)** Results of fine-tuning the human_gpt2-v1 model. The top grey line is the percentage of sequences in different length ranges, the red is the maximum tokens allowed by the model, and the RUN_Vote is the result of three independent runs of voting (RUN0; RUN1; RUN2).

### 3.4 Model Feature Learning Performance in the HERV Dataset

After fine-tuning on the HERV dataset, the advanced pre-trained models DNA_bert_6 and human_gpt2-v1 demonstrated effective sequence classification performance. To gain a deeper understanding of the representation learning ability by fine-tuned large models, we visualized the latent patterns learned through unsupervised learning from the dataset, as well as the representations learned by the models respectively (Fig.4). The DNA sequences in the test set were parsed into 6-mers, yielded high-dimensional sparse matrices which were subsequently dimensionally reduced and visualized using SparsePCA and TruncatedSVD (Fig.4A; Fig.4B). The visualization revealed that specific sequence patterns matching the labels were present in the dataset, HERV and Non-HERV sequences in non-coding regions exhibited clear differences, whereas HERV and Non-HERV sequences in coding regions showed smaller differences and clustered together with the points in non-coding regions. To evaluate the models’ ability to recognize sequence features, we utilized the last hidden layer of the fine-tuned DNA_bert_6 model (RUN0) for the HERV multi-classification dataset. We employed PCA and t-SNE methods to visualize the hidden layer representations, and the results showed that our fine-tuned DNA_bert_6 model successfully learned the patterns specific to HERV sequences and could distinguish HERV sequences within coding regions from HERV Non-coding regions more distinctly. PCA dimensionality reduction clearly captured differences between multiple classification labels, while t-SNE method revealed differences between classification labels and also highlighted subgroups within specific label types(Fig.4C; Fig.4D). We marked the ∼3.92% overlapping HERV sequences in the test set with the HervD_Atlas database as reference points, and further visualization of the t-SNE results indicated that model learned the specific patterns of ERV1, ERV2, and ERV3 subtypes in the HERV sequences through unsupervised learning, and could differentiate them. However, it is crucial to note that it remains unclear whether the model classified subgroups based on the length differences of these three types of HERV sequences [29]; **Supplementary Fig.3**). Furthermore, the last layer feature visualization of the fine-tuned human_gpt2-v1 model also demonstrated that this pre-trained model learned the latent patterns within the HERV dataset (**Supplementary Fig.2**).

**Fig. 4.**
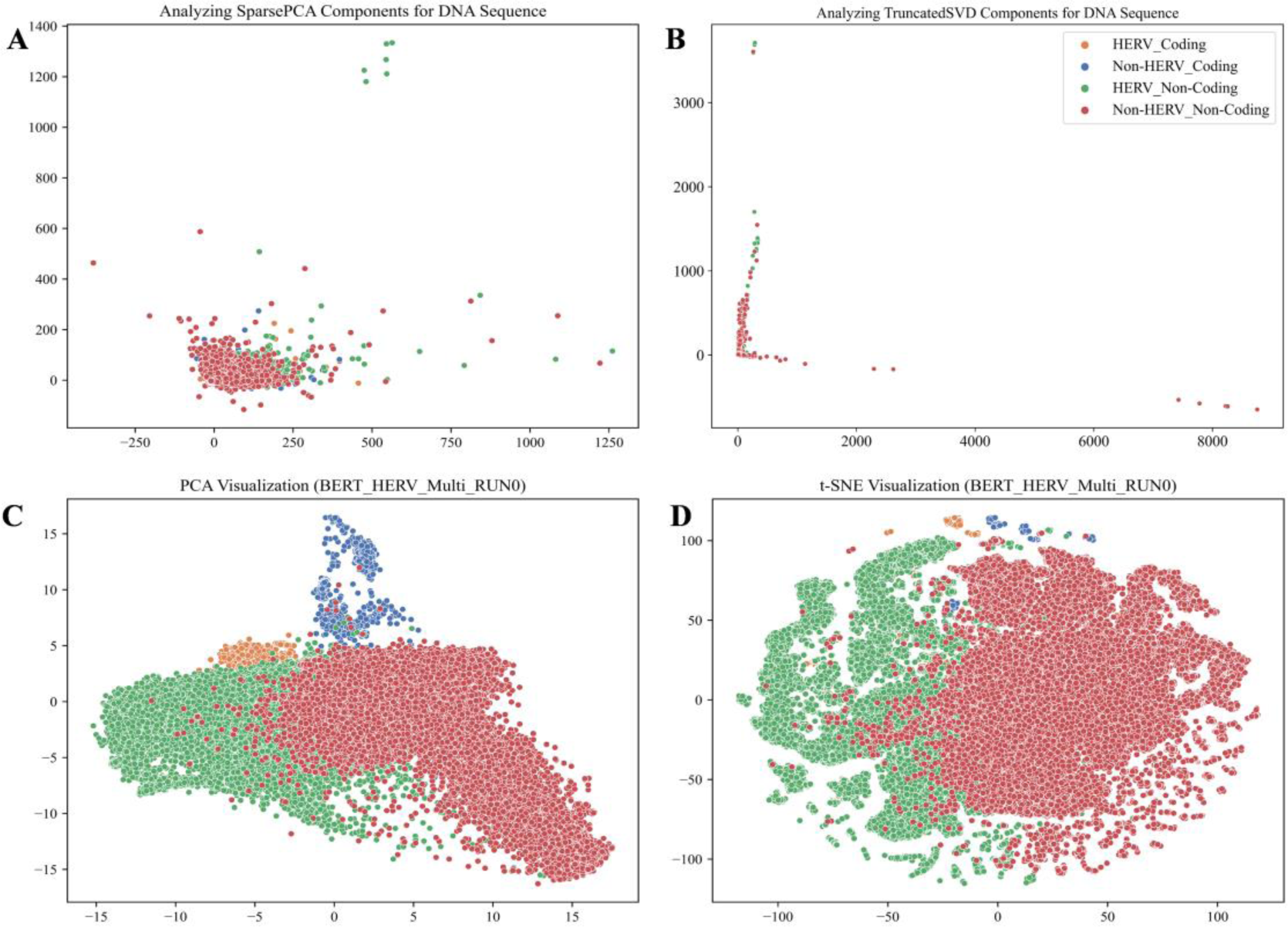
Model representation learning of potential feature patterns in the HERV dataset. **(A)** Visualization of SparsePCA downscaled components of the original DNA sequences; **(B)** Visualization of TruncatedSVD downscaled components of the original DNA sequences; (**C**) Visualization of PCA downscaled components of the last hidden layer in the fine-tuned model; (**D**) Visualization of t-SNE downscaled components of the last hidden layer in the fine-tuned model.

### 3.5 Phenotype-Specific ALRW Scores in the HERV Dataset

In our investigation of the attention scores assigned to different regions within DNA sequences by the DNA_bert_6 model after fine-tuning on the HERV dataset, we extracted the attention score matrices from the final layer of the fine-tuned model on the HERV test dataset. We then calculated the average local representation weighted (ALRW) score, which represents the overall average attention score of phenotype-specific sequences (**Supplementary Fig.4**; Fig.5A). The ALRW scores of different phenotypes in the HERV dataset exhibited variation across the 512 tokens of the input sequences (Figure 5A). All phenotypes showed higher ALRW scores at the beginning and end of the tokens. The ALRW scores of the non-coding HERV sequences (HERV_Non-Coding) were highest at the beginning and end, but lower in other parts compared to other phenotypes. However, the overall distribution of ALRW scores for the coding HERV sequences resembled that of the Non-HERV sequences. Notably, the ALRW scores for the twelve multi-head attention blocks on the non-coding HERV sequences within the test dataset demonstrated a distinct distribution (Supplementary Fig.6). Furthermore, the distinct distributions of ALRW scores within the 12 multi-head attention blocks indicated that different attention heads captured varied feature representations from the sequences (**Supplementary Fig.5**).

**Fig. 5.**
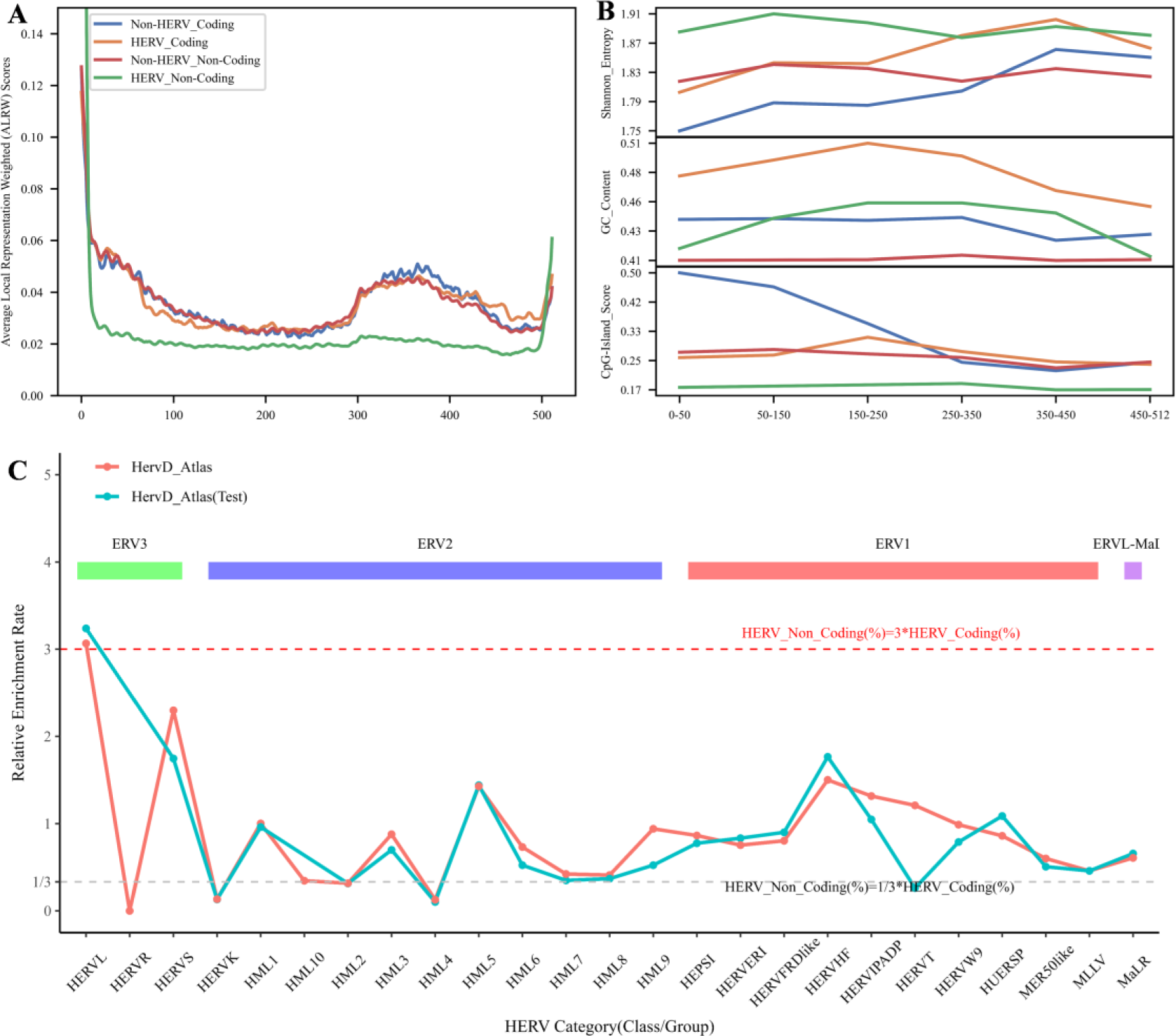
In-depth analysis of average local representation weight scores (ALRW) for specific phenotypic sequences. **(A)** Overall ALRW score distribution of specific phenotypic sequences; **(B)** the characterization of sequences within different tokens positional regions; **(C)** Relative enrichment rates of specific Group sequences within different HERV subtypes (ERV1; ERV2; ERV3; ERVL-ma) in overlapping sequences of the HervD Atlas database.

To further investigate the reasons for the differences in ALRW scores among the various phenotypes, we truncated the HERV phenotypic sequence in the corresponding regions based on token positions. Then, we calculated the GC content, Unique 6-mer frequency, sequence information entropy, and potential CpG island scores of these sequences. It can be observed that the non-coding HERV sequences (HERV_Non-Coding) had higher sequence information entropy and Unique 6-mer frequency, as well as lower CpG island scores, while both the non-coding and coding region HERV sequences showed higher GC content (Fig. 5B; **Supplementary Fig.7**). Moreover, the test set of HERV sequences that could be classified by the HervD_Atlas database revealed that compared to the coding region sequences, the non-coding region sequences had a greater proportion of ERV3-type HERVs and a lower proportion of ERV2-type HERVs (Supplementary Fig. 8). The phylogenetic tree of ERV sequences indicated that ERV3-type sequences were older than ERV1 and ERV2 [29], implying that their prevalence in non-coding regions might result from evolutionary silencing, which is consistent with the finding that the ALRW score distribution is significantly different from other phenotypes. To further explore this phenomenon, we calculated the relative enrichment rates of non-coding versus coding region HERV sequences in the HervD_Atlas dataset (Fig.5C). The results indicated a marked enrichment of ERV3-type HERVL sequences (Solitary LRT) in non-coding regions, characterized by the loss of viral coding capabilities. These sequences are highly conserved and ubiquitously distributed among mammals [30, 31]. Conversely, ERV2-type HERVK sequences, preserving the original provirus structure, were significantly enriched in coding regions. It was also shown that HERVK (HML-2) is not fixed in the gorilla genome but is present in the Neanderthal and Denisovan genomes[30], highlighting its recent positive selection and integration into the human genome.

Based on the characteristics observed in these sequences, we can roughly summarize the potential reasons for the overall low Average Local Representation Weighted Score (ALRW) of the non-coding region HERV sequences as follows: **1) Enhanced complexity and diversity**: The high sequence information entropy and unique 6-mer frequency of non-coding HERV sequences indicate their complexity and diversity, which makes it challenging for the model to capture useful features, resulting in lower scores; **2) Evolutionary silencing and balance**: Non-coding HERV sequences may have been subjected to more extensive silencing events during evolution, leading to differences in their activity, expression, or functionality compared to other sequences, such as low CpG island scores, loss of HERVL viral coding sequences, etc.; **3) A leaning bias towards learning prominent sequence features**: The non-coding HERV region contains more Solitary LRT sequences, and the model tends to focus on assimilating these prominent features, thereby reducing the ability to capture complex features in other HERV sequences.

### 3.6 Motifs Analysis of HERV Dataset with High ALRW Scores

#### 3.6.1 Enrichment and Pathogenicity Analysis of Motifs in High ALRW Score Regions

The research findings suggest that high ALRW scores effectively identify motifs in the test set sequences, which display both specific and shared features across various phenotypes. We identified sixteen types of HERV sequence-specific motifs in both coding and non-coding regions, while unique motifs were also discernible in these regions (Figure 6A). Similar trends were observed when analyzing motifs using the full-length sequences from the test set (**Supplementary Fig.9A**). To assess the specificity of motif enrichment in HERV sequences, a hypergeometric distribution was employed to statistically analyze motifs in both the coding and non-coding regions.

**Fig. 6.**
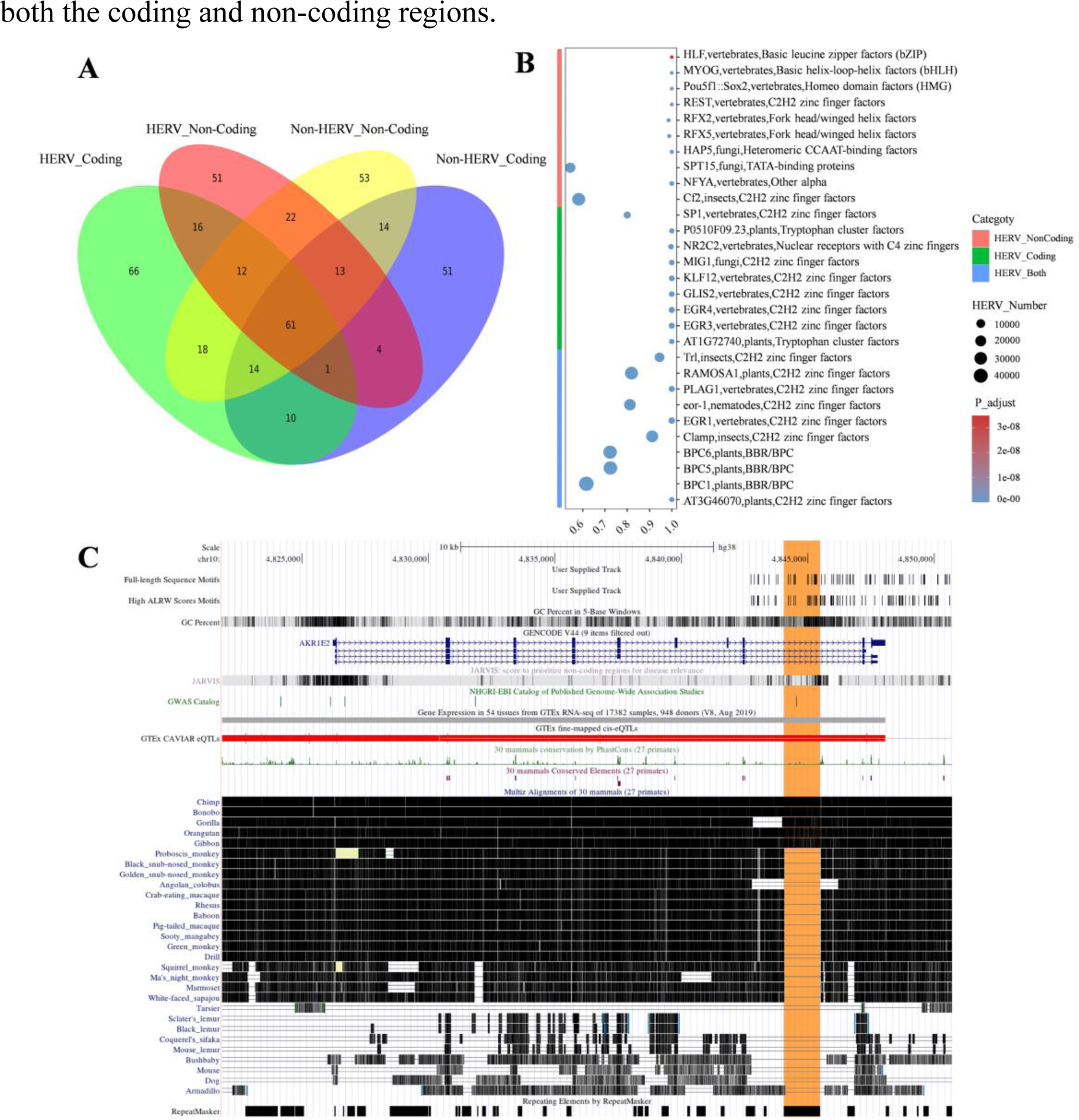
Identification analysis of motifs within phenotype-specific high ALRW scores sequences. **(A)** The overlap of motifs was determined for four specific phenotypes; **(B)** HERV phenotype sequence-specific motifs enrichment analysis demonstrated, The x-axis of the analysis represents the overall rate of HERV type motifs; **(C)** The multiple motifs-enriched genes screened by two strategies, named AKR1E2. Hominoidae-specific HERV sequences are highlighted in orange (chr10_ERV1_01040; Hominoidae: Human; Chimp; Bonobo; Gorilla; Gibbon; chr10:4844040-4845480; ERV1 family ;HERVW9).

The results revealed a significant enrichment of HERV-specific motifs when filtering motifs through high ALRW scores compared to the identification method that uses full-length sequences. This indicates that the high ALRW scores can effectively capture and filter out unique patterns in HERV sequences (**Supplementary Fig.9B; Supplementary Fig.9C**). Upon categorizing the HERV-specific enriched motif sequences identified by high ALRW scores into HERV_Both, HERV_Coding, and HERV_Non-Coding based on their frequencies, motifs associated with DNA-binding and gene-regulation, such as C2H2 zinc finger factors, Tryptophan cluster factors, and AP2/EREBP, were enriched in both the coding and non-coding regions of HERV sequences [32–35]. Conversely, motifs like Nuclear receptors with C4 zinc fingers were predominantly found in coding regions, whereas transcription-related motifs, such as TATA-binding proteins, were abundant in non-coding areas[36, 37] [38, 39](Fig.6B). In summary, specific enriched motifs present in HERV sequences (coding and non-coding regions) are implicated in gene expression regulation, with some unique motifs involved in recognizing nuclear receptors within coding regions. Additionally, motifs identified from the full-length sequences also demonstrated enrichment, with those like Rbpjl, vertebrates, Rel homology region (RHR) factors playing roles in immune regulation, development, and inflammation regulation processes in organisms[40–42]. (**Supplementary Fig.10**).

Using HERV-specific motifs identified by high ALRW scores (coding and non-coding regions), we constructed a HERV sequence-specific non-overlapping motifs interval of 311.857 kb. Within this interval, the major genes overlapping within the motifs interval are primarily located in the Protein_coding and LncRNA regions, involving functional sequence elements such as Enhancer, CTCF_binding_site, and open_chromatin_regio (**Supplementary Fig.11**).We observed multiple specific enrichments of motifs in genes related to neurodevelopment and synaptic function (*NRXN1, CNTNAP2, RBFOX1*), cellular processes and cancer (*FHIT, DNAJB13*), cell adhesion and intercellular communication (*CTNNA3, ANKS1B*), and signal transduction (*BANK1, PDE10A, NKAIN3*). Functional enrichment analysis of all protein-coding genes and disease relevance analysis revealed that genes associated with the specific enriched motifs were involved in processes such as neural activity, cellular morphology and transport, metabolic regulation, and GPCR signaling. Furthermore, they showed close associations with spinal health, neural development and intelligence, and exhibited specific enrichment in DRG neuronal cells (**Supplementary Fig. 12**).

To further investigate the induction of disease by specific enriched motifs, we integrated information from the HervD_Atlas database with the pathogenicity scores predicted by AlphaMissense[26] and PrimateAI[19]. Utilizing the HERV-specific non-redundant motif sequence intervals identified by high ALRW scores, a cumulative length of 47,759 kb (∼15.31%) mainly from the ERV1 family overlapped with sequences in the HervD_Atlas database, which are involved in diseases such as cancer and the nervous system. Compared to the motifs identified based on the full-length sequences, the motifs identified based on high ALRW scores leaned towards high-frequency HERV sequences (average of all specific phenotypic sequences and multi-head attention layers), resulting in the loss of low-frequency sequence information but also the identification of new motifs. **Employing both strategies**, we successfully identified 8 shared motifs (AT3G46070, BPC1, BPC5, BPC6, KLF17, NR2C2, SP2, SP8) in the AKR1E2 gene. Within this gene, there is a Hominoidae-specific HERV sequence chr10_ERV1_01040 exhibited low expressed in 8 cancerous tissues but demonstrated high pathogenicity scores at specific sites (**Supplementary Table 3; Supplementary Table 4; Supplementary Table 5**). The motif sequences AT3G46070, SP8, and KLF17 within this interval play important roles in processes such as cell proliferation and metabolism, brain development[43], and cancer invasion[44](Fig.6C;**Supplementary Fig. 14**). Furthermore, motifs like Eor-1(chr6:28692872-28692886), identified between genes in the chr6_ERV2_01565 (Hominoidae; ERV2 family; HERVK) sequence, which is significantly upregulated in liver cancer (Supplementary Table 3). **Regarding HERV-specific motifs identified through high ALRW scores**, it is worth mentioning that the HERV sequence chr9_ERV2_01861 (Hominoidae; ERV2 family; HML4) contains the CTFC motifs (chr9:76666118-76666151) identified by high ALRW scores within the PRUNE2 gene. This particular HERV sequence is lowly expressed in 4 cancerous tissues and encompasses a CRE regulatory element (**Supplementary Fig. 13A**; **Supplementary Table 3; Supplementary Table 5**). **Lastly, specific motifs identified through full-length sequence analysis,** motifs like ZNF135, which are associated with protein folding and activation processes, have a close correlation with a wide range of biological functions and diseases[45, 46] (**Supplementary Table 4**; **Supplementary Table 5**).

#### 3.6.2 Species Conservation of HERV-Specific Enriched Motif Sequences

Using fine-tuned models to obtain the average local representation weight scores (ALRW) of DNA sequences, our study pinpointed motifs uniquely enriched within human HERV. These HERV-specific motifs sequences are involved in the regulation of essential biological activities in organisms and are closely related to diseases such as the nervous system and tumors. To further evaluate the characteristics of HERV-specific motifs, we investigated their diversity among human populations and their conservation across primate species (Fig.7). By combining published human pan-genomic data, we analyzed the sequence polymorphism (alignment depth) of HERV-specific motifs within their corresponding intervals and visualized them based on overlapping gene types, indicating that LncRNA and protein-coding genes have higher polymorphism (Fig.7A). We also identified highly conserved motif sequences in the CCDC200 gene within the human population, but significant differentiation was observed in primates. It is worth noting that the CCDC family proteins are believed to be involved in various physiological and pathological processes, including gametogenesis, embryonic development, hematopoiesis, angiogenesis, ciliogenesis, and even the mechanisms of cancers[47, 48] (Fig.7B; **Supplementary Fig.15**). Additionally, regions with highly differentiated motifs in the human population or genes containing multiple repetitive segments within the genome are associated with immune function and inflammatory responses (*CD200; CD226; CLEC4E*), signal transduction, cell communication, and the nervous system (*ADCY1; ABL1; ADRA1A; NRG3; RAB3GAP1; OTUD7A; CSGALNACT2*), and metabolic processes (*CYP27A1; AACS; ALDH3A2*). This suggests their role in in the transcriptional regulation of divergent traits within primates and their co-evolution with pathogens in the environment [30, 49, 50](Fig.7). Moreover, motifs conserved among primates that exhibit a degree of polymorphism in the human population may be involved in the development of human organs and the adaptability of the immune system to the environment. By comparing brain organ samples from humans and great apes, it was found that the epithelial-mesenchymal transition regulator ZEB2 promotes the transformation of neural epithelium, which ultimately leads to the expansion of the human brain[51].The membrane receptor ROR1, which is crucial for embryonic development and overexpressed in multiple cancers, has been shown to be a safe and practical target for CAR-T cell immunotherapy in clinical trials using non-human primates[52, 53].

**Fig. 7.**
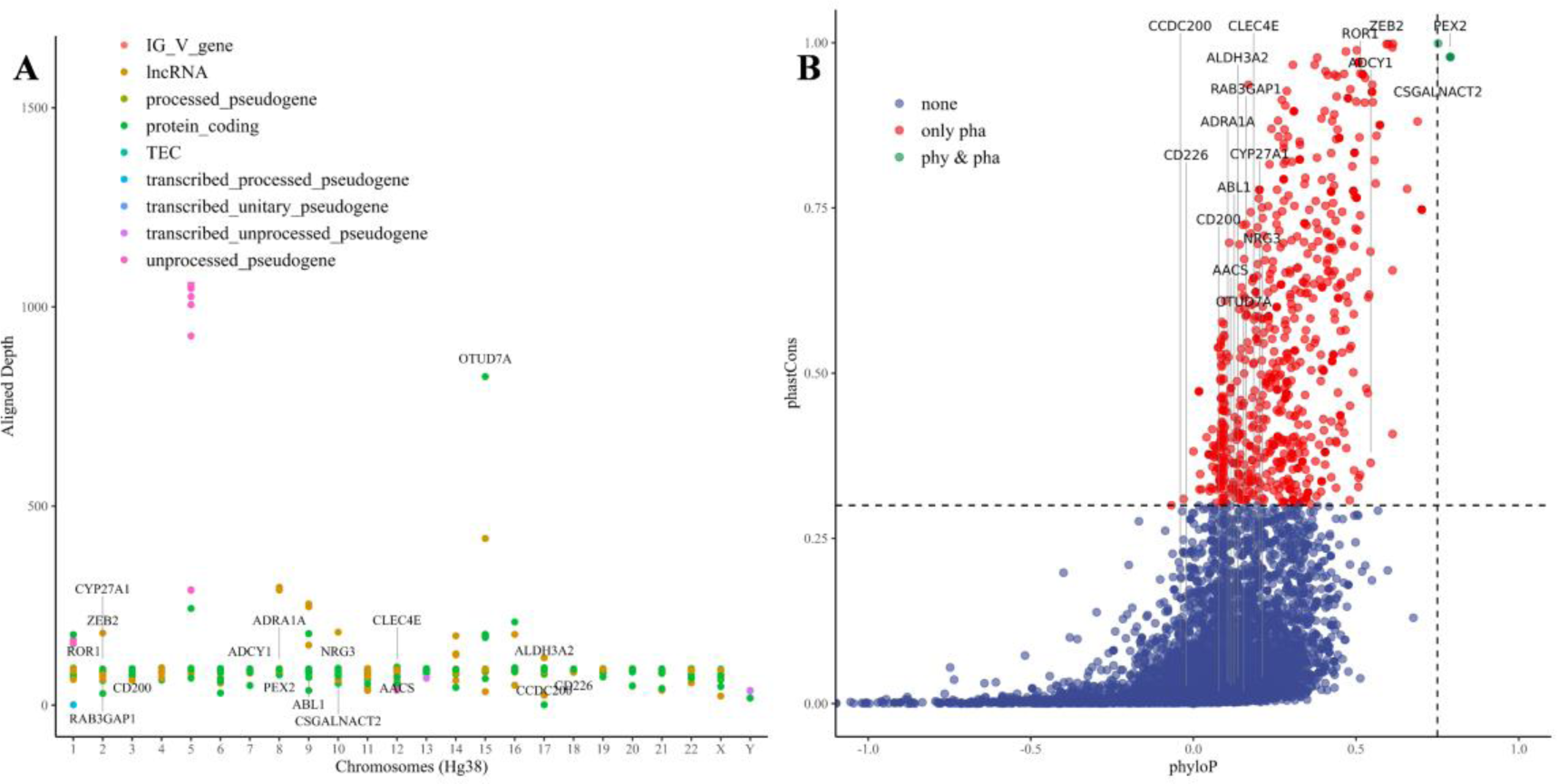
Conservation of species within phenotype-specific enriched motifs regions. **(A)** Pan-genomic representation of population polymorphisms within phenotype-specific enriched motifs regions; **(B)** The conservation of these motifs is investigated in primates.

## 4. Discussion

In the post-genomic era, the functional analysis of DNA sequences has paramount importance. We constructed multiple genotype-phenotype dataset (Individual and molecular phenotypes) by integrating accumulated data and knowledge. After fine-tuning the large model with DNA_bert_6 and human_gpt2-v1, we achieved balanced binary and imbalanced multiclass phenotypic classifications with exceptional efficacy. Notably, the fine-tuning on the HERV dataset not only demonstrated enhanced capability in identifying and delineating various informational categories within DNA sequences, but also pinpointed specific motifs associated with neurological disorders and cancers in the region of high local representation weight (ALRW) scores. The conserved analyses of these motifs further revealed the species’ environmental adaptation and its co-evolutionary dynamics with pathogens. These sequence-specific motifs could revolutionize nucleic acid vaccine development and targeted therapeutics for genomic diseases, such as liver cancer intervention vaccine and drug development based on the expanded sequence of the Eor-1 motif within HERVK.

The main challenges of DNA sequence analysis include the complex sequence features and model input length limitations. DNA sequences usually encapsulate vast information and complicated structures, such as repetitive sequences, etc., making it difficult to extract useful knowledge using traditional methods. Moreover, variability in the loss function outcomes across sequences of varying complexities sequence, which may require model pre-training by sequence type. In addition, the population biases and sequence frequency may generate learning bias in the model and affect downstream tasks. Currently, the commonly used pre-training BERT and GPT models have a maximum model input tokens limitation, possibly resulting in loss of spatial information of the genome and important regulatory elements, such as the long-distance Enhancer. Despite DNA controlling complex life activities, research predominantly focuses on approximately 3% of protein-coding sequences in the genome. Pre-trained genome models can improve understanding of the DNA genetic blueprint, which is vital for deciphering gene control of biological traits and disease development.

Future research will focus on developing incremental pre-training models for the human pangenome, incorporating model knowledge distillation, and integrating knowledge graphs, etc. We will also explore innovative frameworks capable of handling extended input tokens, such as HyenaDNA and LongNet [9, 54]. Ultimately, it is expected that our pangenome incremental pre-training model can match the capabilities of the reference sequence hg38, potentially transforming the computational biology landscape. This advancement in genome-level representation learning will deepen our understanding of life’s nature and accelerate the progress in natural language processing technologies. Furthermore, the findings and data from this research will contribute to personalized therapy strategies, including vaccine and drug development targeting HERVK sequences[55].

## 5. Conclusion

This study employs a pre-trained genome model to identify and interpret genetic signals associated with specific phenotypes. The findings can be categorized into three main aspects:**1) Genotype-Phenotype Dataset Aspect**: We utilized various genotype-phenotype datasets to assess the effectiveness of fine-tuned large models in balanced binary classification and imbalanced multi-class sequence challenges. Using the HERV dataset, we evaluated the variation in model classification metrics across different sequence length distributions, exploring the correlation between data distribution patterns and model performance. These datasets play a critical role in assessing the efficacy of pre-trained genome models and they highlight the importance of model selection based on data distribution, requirements, and costs. **2) Representation Learning in Genome Pre-trained Models**: The fine-tuned HERV dataset reveals that hidden layer features enable the model to recognize phenotypic information in sequences and reduce noise. To investigate how the model isolates phenotypic label-specific signals, we calculated local representation weight scores (ALRW) for phenotypic labels using average attention matrices. The distribution of ALRW scores aligns with the fundamental characteristics of coding and non-coding areas in HERV sequences and their evolutionary subtypes, validating the utility of genome pre-trained models in the deep analysis of biological sequences. **3) New attempts to integrating Genome Pre-trained Models with Classical Omics Analysis**: By selecting sequences with high ALRW scores specific to phenotypes for motif enrichment analysis, we identified HERV-specific motifs that are implicated in neurological diseases, tumors, and other biological processes.Therefore, they have potential applications in vaccine and targeted drug discovery. Furthermore, the polymorphism of these motifs in human populations and their conservation in primates provide insights into primate adaptation to environmental pressures and integration of pathogens into the host genome.This insight aids in selecting motifs for further research and development in vaccines and drugs, using non-human primates, exemplified by HERVK sequences.

## Author Contributions

Conceptualization: DD, FZ, LL; Formal Analysis: DD; Funding Acquisition: LL; Investigation: DD; Methodology: DD; Project administration: FZ, LL; Resources: LL; Supervision: FZ, LL; Visualization: DD; Writing – original draft: DD; Writing – review & editing: DD, FZ, LL.

## Declaration of generative AI in scientific writing

AI-assisted technologies were employed to augment readability and language proficiency. Nevertheless, the final results were exclusively produced by the authors, who meticulously edited the language to conform to domain terminology. Consequently, we are wholly responsible and accountable for the content of this study.

## Acknowledgements

We acknowledge the support provided by the project of Shanghai’s Double First-Class University Construction, the Development of High-Level Local Universities: Intelligent Medicine Emerging Interdisciplinary Cultivation Project, and Medical Science Data Center of Fudan University.

## Code and Result Availability

The code for this project can be accessed at https://github.com/GeorgeBGM/Genome_Fine-Tuning and the relevant key result files related to this topic are accessible for download via 10.5072/zenodo.5983.

## Appendix A. Supplementary data

Supplementary material related to this article can be found online at XXXXXX.

